# *De novo* variants in Chinese ASD trios reveal genetic basis underlying autism without developmental delay and intellectual disabilities

**DOI:** 10.1101/2021.09.30.462503

**Authors:** Jincheng Wang, Juehua Yu, Mengdi Wang, Lingli Zhang, Kan Yang, Xiujuan Du, Jinyu Wu, Xiaoqun Wang, Fei Li, Zilong Qiu

## Abstract

Autism spectrum disorder (ASD) is a complex neurodevelopmental disorder that causes a range of social communication and behavioral impairments. ASD typically manifests in young children, often with developmental delay or intellectual disabilities (DD/ID) as comorbidities. Accruing evidence indicates that ASD is highly heritable and genomewide studies on ASD cohorts have defined numerous genetic contributors. Notably, most of these studies have been performed with individuals of European and Hispanic ancestry and thus there is a paucity of genetic analyses of ASD in the East Asian population. Here, we performed whole-exome sequencing on 772 ASD trios from China, combining with a previous study of 369 Chinese ASD trios, to identify *de novo* variants in a total of 1141 ASD trios. We found that ASD probands without DD/ID carried less disruptive *de novo* variants, including protein-truncating and missense variants, than ASD with DD/ID. Surprisingly, we showed that expression of genes with *de novo* variants found in ASD probands without DD/ID were enriched in a specific group of neural progenitor cells, suggesting a potential mechanism underlying high-functioning autism. Importantly, some ASD risk genes from this study are not present in the current ASD gene database, suggesting that there are novel genetic contributors to ASD in East Asian populations. We validated one such novel ASD risk gene – *SLC35G1* by showing that mice harboring heterozygous deletion of *Slc35g1* exhibited defects in social interaction behaviors. Together, this work nominates novel ASD risk genes and indicates that ASD genetic studies in different geographic populations are essential to reveal the comprehensive genetic architecture of ASD.

## Main text

Autism spectrum disorders (ASD) refers to a collection of neurodevelopmental disorders, mostly occurred before 3 years old or during the toddler stage, associated with impairments in social communication and prominent repetitive behaviors^1^. ASD appears to be a highly heritable disorders^2^. Indeed, harnessing the rapid progress of whole-exome sequencing (WES) and whole-genome sequencing (WGS) technologies, researchers have performed WES and WGS on large ASD cohorts and identified *de novo* and rare inherited variations are major contributors to ASD^3,4^. Moreover, analysis of simplex families (two unaffected parents, one child with ASD, and unaffected siblings), has revealed that *de novo* likely gene-disrupting variants (LGDs), usually causing protein truncations, occur more frequently in children with ASD than in their unaffected siblings^4^.

Although developmental delay or intellectual disability (DD/ID) are not core ASD symptoms, a substantial portion of ASD patients exhibit signs of DD/ID (ID diagnosed after 4 years old). In a recent ASD genetic study containing the largest cohort thus far, the authors classified the ASD risk genes into two categories, ASD-predominant (ASD_p_), genes which are primarily associated with core symptoms of ASD and ASD-neurodevelopmental delay (NDD) (ASD_NDD_) genes which exhibited mutations in ASD probands with severe neurodevelopmental symptoms^5^. Although whether there are “autism-specific” genes is still under debate, a consensus has emerged investigating on genes mainly affecting social functions, instead of general cognitive functions, may shed light on the neurobiology and genomics of social behaviors of animals^6^.

In this study, we performed WES on a Chinese cohort containing 772 ASD probands with both neurotypical parents, in which there are 334 probands showed intelligence quotient or developmental quotient (IQ/DQ) over 70 (referred to as High-functioning autism: HFA, also known as ASD without DD/ID), and 415 probands with IQ/DQ less than 70 (referred to as Low-functioning autism: LFA, also known as ASD with DD/ID). By comparing the component of *de novo* variants identified between HFA and LFA, we found that there are significantly more damaging *de novo* LGD and missense variant happening in LFA probands, comparing to HFA probands. Numerous new ASD candidate genes emerged, including *SLC35G1*, in which we identified recurrent mutations in HFA probands. We provide evidence that it plays a critical role in regulating social behavior in mice because heterozygous *Slc35g1* haploinsufficient mice exhibit deficits in social interaction behaviors.

### Distinct *de novo* variants in LFA and HFA probands

We first analyzed WES of 772 ASD trios (probands and both unaffected parents) collected from the Department of Developmental Children Care, Xinhua Hospital Affiliated with Shanghai Jiao Tong University School of Medicine. We focused on *de novo* variants (present in probands but not in parents) in this study. We identified 401 *de novo* synonymous variants and 708 rare *de novo* non-synonymous variants (allele frequency < 0.1% in the four genomic databases: 1000G-ALL, 1000G-EAS, ExAc-ALL, ExAc-EAS, database websites see Methods, Fig. 1a, Fig. S1). Among 708 rare *de novo* non-synonymous variants in protein-coding regions, there were numerous LGD and missense variants, of which we validated 89.55% of total rare *de novo* non-synonymous variants by Sanger sequencing (Fig. 1a, Table S1).

**Figure 1.**
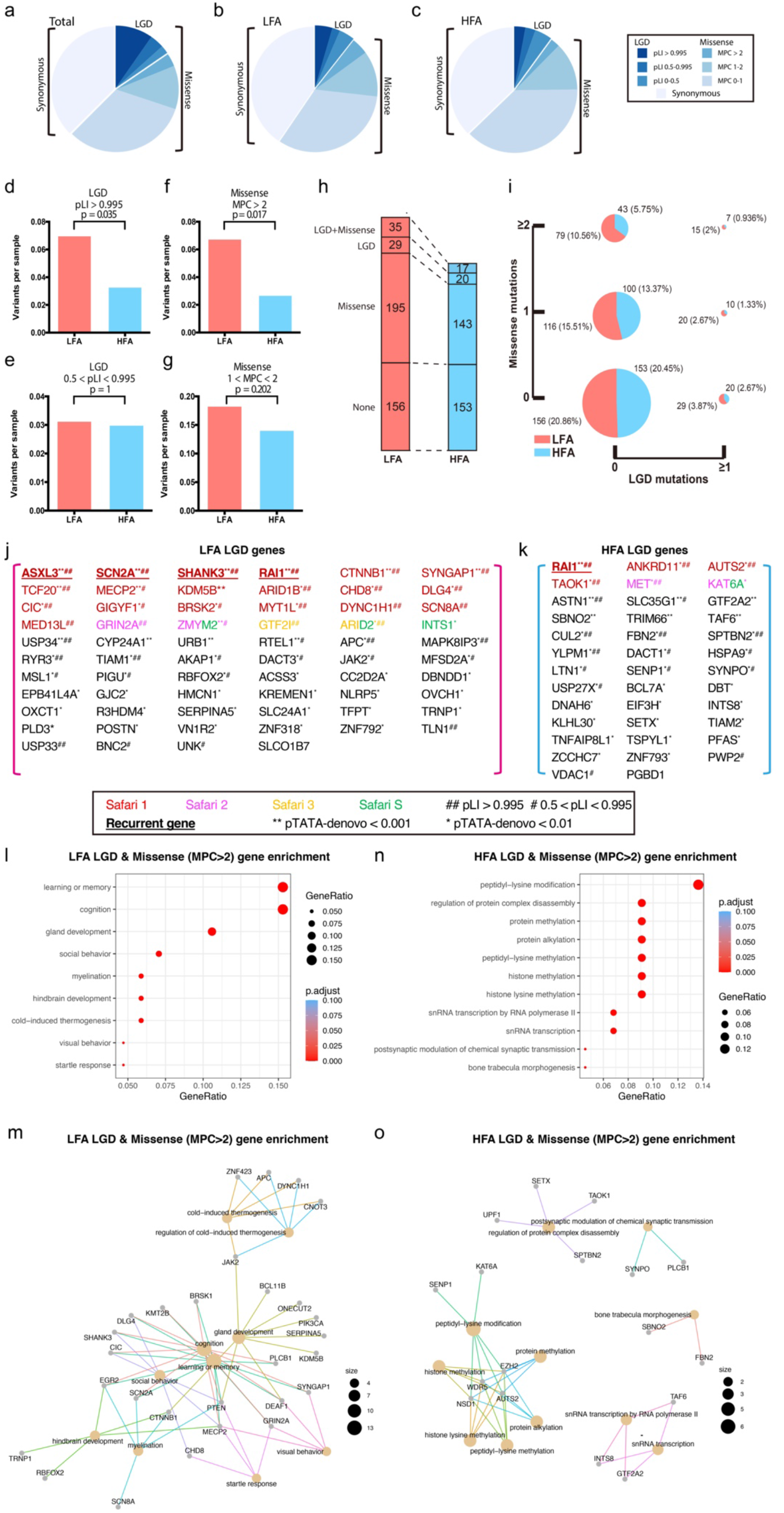
Distribution of LGD and missense variants in LFA and HFA probands. (a-c) The proportion of rare autosomal *de novo* variants split by predicted functional consequences, represented by color, is displayed for Total (a), LFA (b) and HFA (c) probands. LGD and missense variants are split into three tiers of predicted functional severity, represented by shade, based on the pLI and MPC metrics, respectively. (d-g) The variant frequency in *de novo* LGD and missense between LFA and HFA probands based on pLI and MPC metrics. (h) Illustration of numbers of LFA or HFA probands with detection of *de novo* missense variants, LGD variants or both. (i) Percentage of LFA and HFA probands with *de novo* LGD or missense variants. Summary of *de novo* LGD variants identified in LFA (j) and HFA (k) probands. Colors (red, magenta, yellow and green) denote the SFARI genes in category 1, 2, 3, and S, respectively. pTATA-denovo value (** < 0.001, * < 0.01); pLI value (## > 0.995, # >0.5 & <0.995); genes with *de novo* recurrent LGD genes were shown in bold and underline. Dot plots (l & n) and cnetplots (m & o) depict enrichment ratios for GO term categories and enriched biological processes of genes with *de novo* LGD and missense (MPC > 2) variants from LFA (l & m) and HFA (n & o) probands.

To classify the severity of protein-changing *de novo* variants, we analyzed the “probability of loss-of-function intolerance” (pLI) score^7^ and the integrated “missense badness, PolyPhen-2, constraint” (MPC) score^8^ for LGD and missense variants, respectively (Fig. 1a-c). Previously, ASD probands were found to carry significantly more LGD variants, but not missense variants, than their unaffected siblings in simplex families^4^. Interestingly, we found that LFA probands carried significantly more both LGD and missense variants likely having larger effects (pLI > 0.995 or MPC > 2) than HFA probands (Fig. 1b-g), suggesting that LFA carried heavier mutation burdens than HFA. Therefore, both *de novo* LGD and missense variants may contribute to the etiology of ASD with development delay.

Overall, we identified *de novo* rare non-synonymous variants (including LGD and missense) in 62% (259 of 415) of families having LFA probands, and 54% (180 of 334) of families having HFA probands (Fig. 1h). Next, to further analyze mutation burdens of LFA and HFA, we examined the *de novo* variants occurred in each proband. We found that there were significantly more LFA probands carrying more than 1 *de novo* variants (LGD or missense) than HFA probands, suggesting that comorbid DD/ID is likely associated with heavier mutation burdens (Fig. 1i).

We analyzed the statistical significance using the TADA-denovo method ^7,9^. Among the genes with *de novo* LGD variants, we found that 24 out of total 64 genes in LFA probands were presented in the SFARI database (http://gene.sfari.org) which is the most comprehensive ASD gene database (Fig. 1j). Interestingly, in the genes with *de novo* LGD variants in HFA probands, there are only 6 of 38 genes were previously reported in the SFARI database (Fig. 1k). Many of these novel ASD risk genes passed the TADA-denovo test (*p* < 0.001 - 0.01), suggesting that they may represent East Asian specific ASD risk genes (Fig. 1j, 1k).

To further investigate the function of genes carrying variants with large effects (LGD or missense MPC >2) discovered in LFA and HFA probands, we performed Gene Ontology analysis and found that genes in LFA probands with likely disruptive variants are primarily associated with learning memory and brain development (Fig. 1l, genes elaborated in Fig. 1m), whereas genes in HFA probands with large effects are mainly associated with various post-translational protein modification, histone modification, and small RNA transcription (Fig. 1n, 1o). These data suggest that the etiology leading to LFA and HFA may be distinct in the course of brain development.

### LFA and HFA genes exhibited distinct expression landscapes

To determine the expression profiles of genes with *de novo* variants in LFA and HFA probands, we first examined the expression of candidate genes in three critical developmental stages, including early embryonic stage (1^st^ - 2^nd^ trimester), late embryonic and early postnatal stage (3^rd^ trimester and infant-toddler), and childhood, adolescence and adult stage. We categorized genes with *de novo* variants into these three groups according to the stage at which they were expressed at the highest level (Fig. S2a-c). We first found that genes with *de novo* LGD variants in HFA probands appeared to contain more early embryonic development genes than genes with *de novo* LGD variants in LFA probands (Fig. 2a)

**Figure 2.**
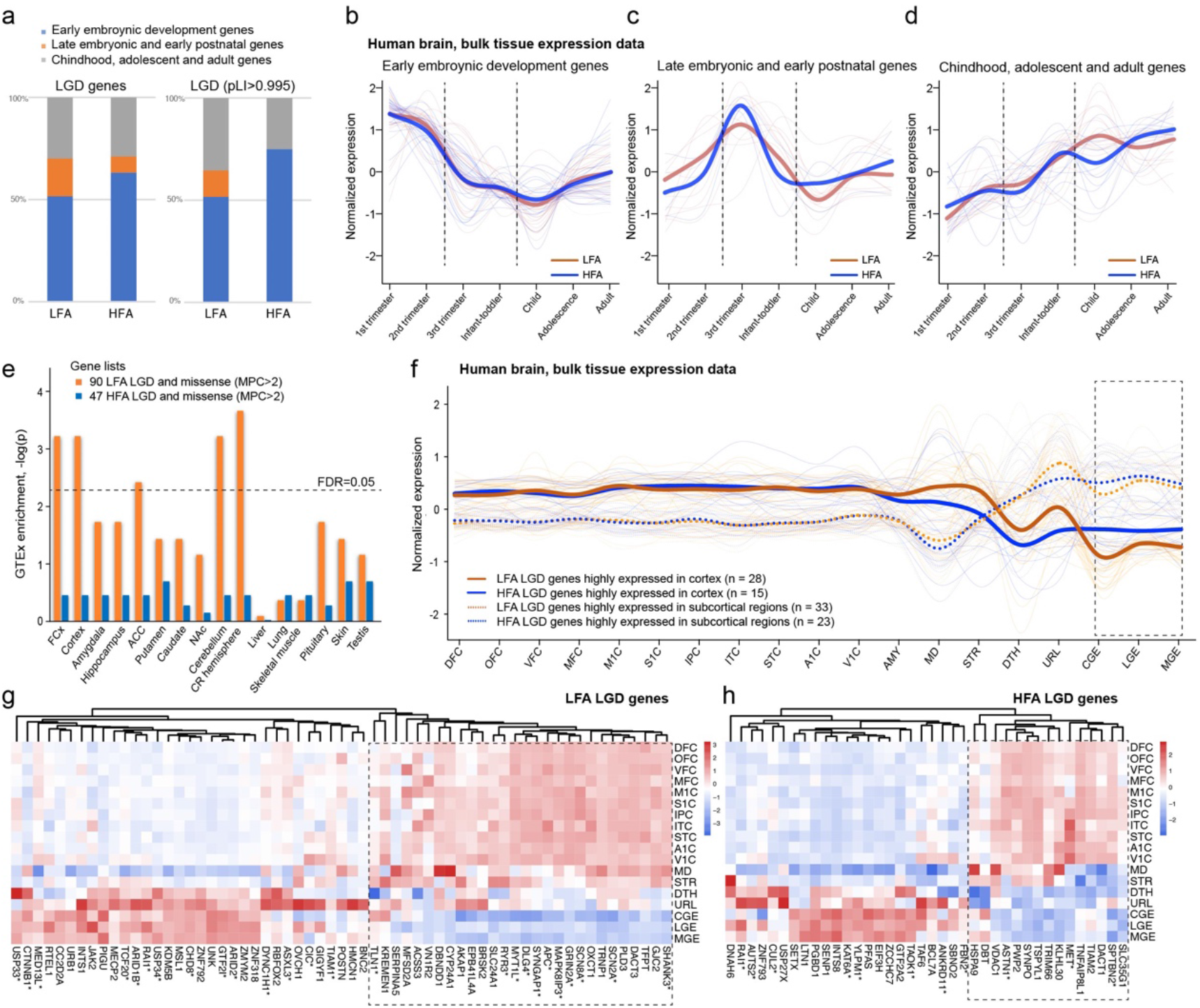
Analysis of LFA versus HFA Genes in the Context of Gene Expression Data. (a) Genes with *de novo* LGD variants were categorized into three groups, including early embryonic development genes, late embryonic development and early postnatal genes, and childhood, adolescent, and adult genes. A percentage stacked barplot shows the proportion of each category of LFA and HFA genes. BrainSpan bulk RNA-seq data was used to plot the normalized expression of the 61 LFA and 38 HFA LGD genes across development, splitted by three stages, including (b) early embryonic development genes, (c) late embryonic development and early postnatal genes, and (d) child, adolescent, and adult genes. (e) GTEx bulk RNA-seq data were processed to identify genes enriched in specific tissues. Gene set enrichment was performed for *de novo* LGD and missense (MPC > 2) variants in 90 LFA and 47 HFA genes for each tissue. Sixteen representative tissues are shown, including 10 region-specific brain tissues. (f) BrainSpan bulk RNA-seq data was used to plot the normalized expression of the 61 LFA and 38 HFA LGD genes across 18 region-specific brain tissues, including 11 cortical and 7 subcortical regions. Genes highly expressed in cortex are shown in solid lines, while genes highly expressed in subcortical regions are shown in dash lines. Heatmaps showing BrainSpan bulk RNA-seq data of 61 LFA (g) and 38 HFA (h) LGD genes across 18 region-specific brain tissues. * pLI value > 0.995.

From the gene expression data from human brain in the BrainSpan database, we found that expression levels of genes with *de novo* LGD variants in LFA (n = 61, excluding three genes showing low expression level in the brain: *NLRP5*, *R3HDM4*, *SLCO1B7*) showed a similar pattern with genes with *de novo* LGD variants in HFA (n = 38) at the early embryonic stage (Fig. 2b). Interestingly, genes with *de novo* LGD variants in HFA exhibited higher expression in late embryonic stage and lower expression in early postnatal stages (Fig. 2c). Moreover, genes with *de novo* LGD variants in LFA showed higher expression levels in the childhood stage than genes with *de novo* LGD variants in HFA (Fig. 2d).

We next evaluated enrichment of genes with *de novo* variants of large effects (LGD, missense MPC > 2) with bulk RNA-seq data in the Genotype-Tissue Expression (GTEx) resource^10^. We observed significant enrichment of LFA genes in various brain tissues including frontal cortex, cerebellum (FDR < 0.05) and amygdala, hippocampus, ACC, basal ganglia, nucleus accumbens (*p* < 0.05) (Fig. 2e), suggesting that etiology of LFA may be widely associated with various brain subregions. We further evaluated expression levels of genes with LGD variants in LFA and HFA probands. We found that according to the normalized expression level, genes with LGD could be divided into two groups, a cortically expressed group and a sub-cortically expressed group (Fig. 2f). Interestingly, we found that the cortically expressed groups of HFA probands exhibited higher expression level in central, medial and lateral ganglionic eminence (CGE, MEG, LGE) where the GABAergic neurons are mainly formed prenatally (Fig. 2f-h). This observation suggested that the etiology of HFA may be associated with dysfunction of GABAergic neurons in the brain.

To further examine the expression patterns of genes with *de novo* variants in LFA and HFA at the single-cell resolution, we collected total 17434 transcriptomes from gestational week (GW) 09-26 of human fetus brains^11,12^ and further classified them into sub-types according to marker genes (neural progenitor cells: *PAX6*, glutamatergic/excitatory cells: *NEUROD2*, *SLC16A6*, *SLC16A7*, GABAergic/inhibitory cells: *DLX2*, *GAD1*) (Fig. 3a, 3b).

**Figure 3.**
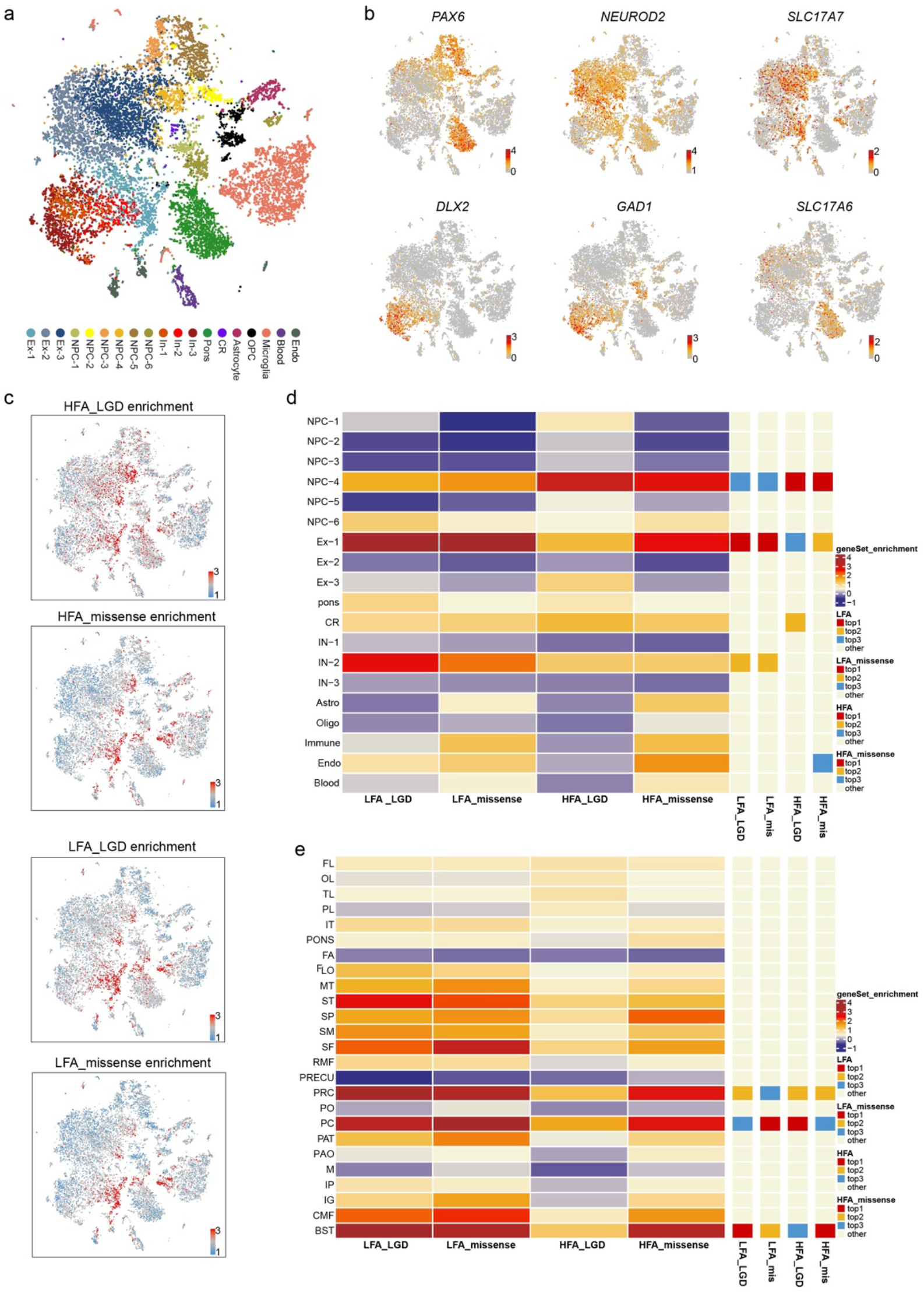
Single-cell expression pattern of *de novo v*ariants in LFA and HFA probands. (a) t-SNE displaying diverse cell subtypes, cells were colored by subtypes. (b) Different population of cells identified by distinct markers. Neural progenitors: *PAX6*, excitatory neurons: *NEUROD2, SLC17A7* and *SLC17A6*, and GABAergic neurons: *DLX2* and *GAD1*. (c) t-SNE showing the enrichment score of genes with *de novo* variants in LFA and HFA probands (high, red; low, blue). (d) Heatmap showing averaged enrichment score of genes with *de novo* variants in LFA and HFA probands among different cell subtypes (high, red; low, blue). The top three most enriched cell types for each gene-set were shown on the right sidebar. (e) Heatmap showing averaged enrichment score of genes with *de novo* variants in LFA and HFA probands among different brain regions (high, red; low, blue). The top three most enriched brain regions for each gene-set were shown on the sidebar. (Abbreviations are listed in Table S2).

Interestingly, we found that expression of genes with *de novo* variants (including LGD and missense variants) in LFA probands showed enrichment in sub-types of excitatory cells (Ex-1) and inhibitory cells (IN-2), whereas genes with *de novo* variants in HFA specifically enriched in a sub-types of neural progenitor cells (NPC-4) (Fig. 3c, 3d), which was previously reported to be a transient state of neural progenitors that expressed neural stem cell genes *HES1* and *VIM*, intermediate progenitor cell genes *EOMES* and *PPP1R17*, neuronal genes *STMN2*, *NEUROD1/2/6* and also the reported layer V gene *NPR3* (Fig. 3c, 3d, Fig. S3a-b) ^12^.

From regional aspects, we found that genes with *de novo* variants in LFA probands exhibited enrichment in precentral cortex (PRC), postcentral cortex (PC) and banks of superior temporal cortex (BST) regions (Fig. 3e, Table S2), which was consistent with our previous report ^13^ and suggested that dysfunction of primary somatosensory and motor cortex may be associated with LFA probands.

### Gene discovery in Chinese ASD cohorts

In order to comprehensively evaluate the contribution of genes with *de novo* variants discovered in the Chinese ASD population, we combined the current cohort with a previous cohort containing 369 Chinese ASD probands with their parents^13^ and analyzed all genes with *de novo* variants in the dataset of 1141 ASD probands.

We identified 59 genes at the false discovery rate (FDR) < 0.3 with the TADA-based model, of which 22 passed FDR < 0.1 and 13 passed FDR < 0.05^9^ (Fig. 4a, Table S3). Among ASD risk genes at the FDR threshold of 0.1 or less, 40 % (9 of 22) are not present in the SFARI gene set, suggesting that a substantial amount of ASD risk genes in Chinese ASD cohorts may be distinct from genes found in other populations previously studied. These new ASD risk genes from Chinese cohorts are enriched for functions in gene expression regulation, neural communication, cytoskeleton & endomembrane system and others (Fig. 4b).

**Figure 4.**
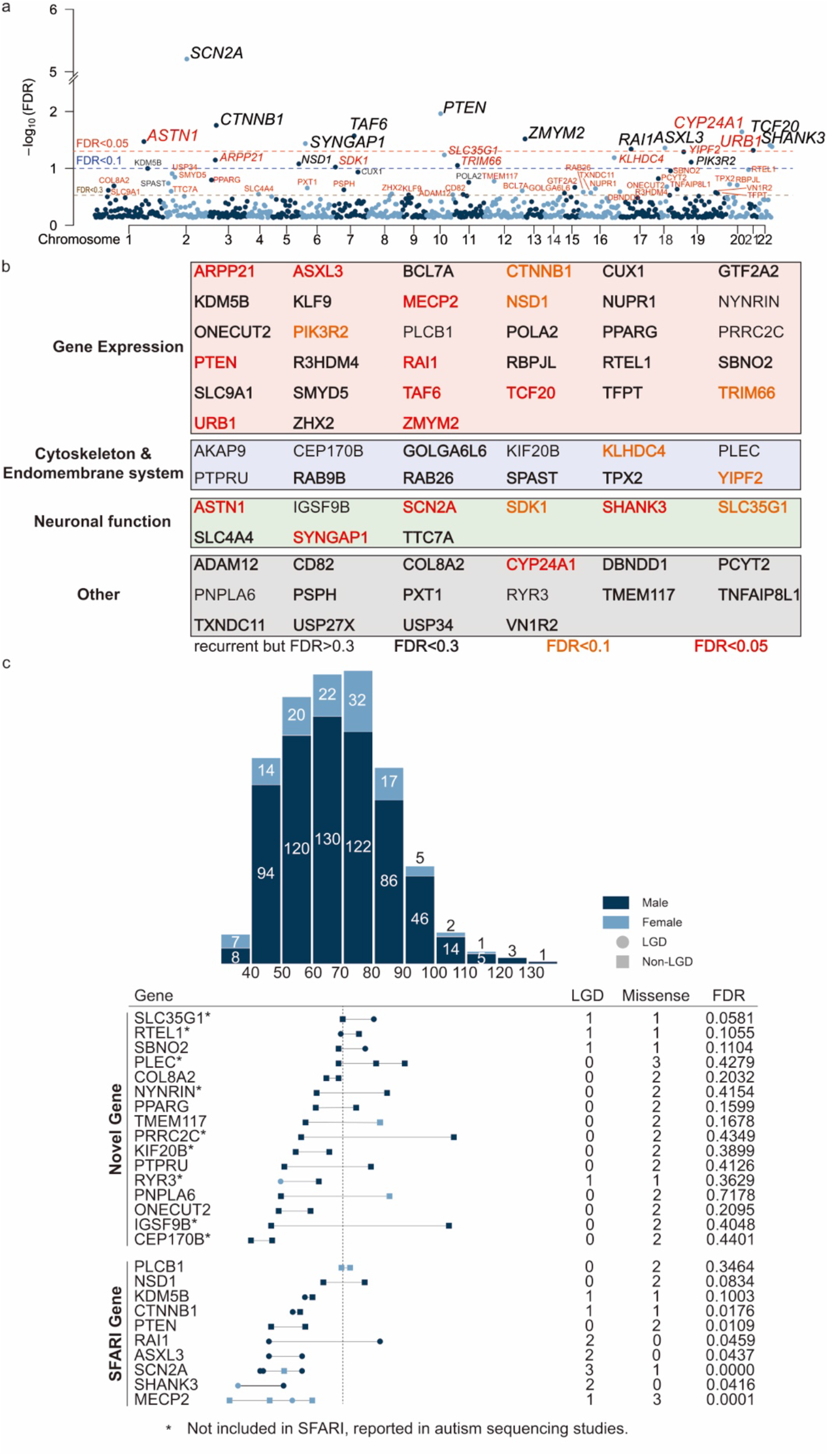
Gene Discovery in the ASD Cohort. (a) Manhattan plot of genes in the 1141 ASD trios analyzed with the TADA model. The model identifies 59 genes associated with ASD at a false discovery rate (FDR) threshold of 0.3 or less. 13 genes pass the threshold of FDR < 0.05, and 22 genes pass the threshold of FDR<0.1. Each point representing a gene. X axis presents chromosome, y axis presents the FDR rate. Genes with black color means they are present in the SFARI gene set, genes with red color stands for potential novel genes which are not collected by SFARI yet. (b) Table of functional groups of genes presented in (a) and genes with recurrent *de novo* variants but with FDR > 0.3. Each group were presented with various color indicated. (c) The IQ/DQ distribution of ASD proband carrying *de novo* variants (upper panel). Affected males (dark blue) account for 83.98%, with mean IQ of 67.97. The mean IQ of affected females (light blue) is 66.88. The distribution of IQ/DQ of ASD probands carrying recurrent *de novo* variants was shown in the lower panel. The vertical dashed line indicates an IQ/DQ of 70 (lower panel). FDR of SFARI gene and potential novel genes were indicated.

To determine if *de novo* variants may be associated with IQ/DQ of probands, we first plotted the distribution of ASD probands according to the IQ/DQ range (Fig. 4c upper panel). Then we listed ASD probands carrying genes with recurrent *de novo* variants along the IQ/DQ axis (Fig. 4c lower panel). Interestingly, among numerous genes with recurrent *de novo* variants, some of them appeared only in probands with normal IQ/DQ (*SLC35G1*, *RTEL1*, *SBNO2*, *PLEC*, *PLCB1*) (Fig. 4c), suggesting that they are candidate risk genes for ASD without DD/ID. Because of differences in genetic penetrance, variants in the same gene may lead to various IQ/DQ in different individuals, such as *NSD1* and *RAI1* (Fig. 4c). This observation is consistent with previous discussion that it may be difficult to define the “autism-specific” genes since different variants in the same gene may lead to dramatically different symptoms including with or without DD/ID^6^. Thus, we took advantage of the opportunity to identify genes primarily contributing to ASD without DD/ID, by focusing on genes with genetic variants that will not lead to DD/ID symptoms in recurrent cases.

### *SLC35G1* is a novel candidate gene for ASD without DD/ID

We identified a novel ASD risk gene *SLC35G1* with the high confidence (FDR = 0.058), which encodes a transmembrane protein transporting nucleotide sugar^14^. We identified two mutations, K120fs and A310P in two unrelated male ASD patients without DD/ID (Fig. 4c, Fig. 5a, 5b). The K120fs mutation is a frame-shifting variant caused by a deletion of 4 base pair (bp), in which 2 bp are located at the end of exon 1, and 2 bp located at beginning of intron 1 of the *SLC35G1* gene. The K120fs mutant leads to premature termination at the 156^th^ amino acid of the SLC35G1 protein. The A310P mutation was predicted as damaging by a series bioinformatic databases including SIFT, Polyphen2, MutationTaster, PROVEAN and fathmmxMKL^15–19^. The alanine at position 310 of the SLC35G1 protein is conserved among zebrafish, xenopus, chicken, mouse, dog, monkey and human (Fig. 5a). It was previously reported that a variant at a splicing site in the *SLC35G1* gene may be associated with ASD^20^. Moreover, the expression of *Slc35g1* enriched in the human brain according to the single-cell sequencing analysis (Fig. S4a, b). Therefore, we speculate that *loss-of-function* variants of *SLC35G1* may lead to ASD.

**Figure 5.**
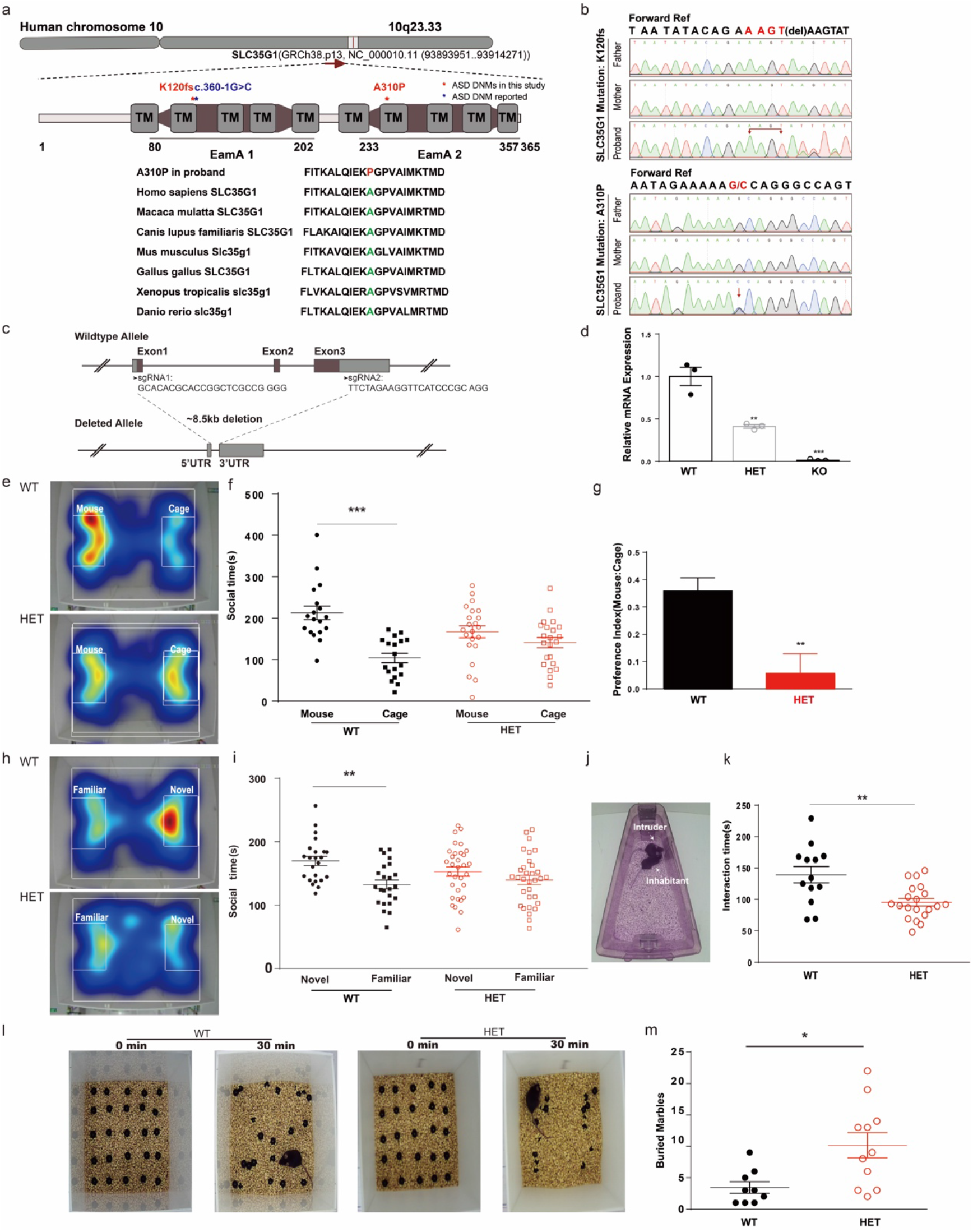
SLC35G1 is a novel risk gene for ASD without DD/ID. (a) Genomic localization of *SLC35G1* (upper panel), three genetic variants found in *SLC35G1* among ASD cohorts (middle panel) and conversation of A310 in various species (lower panel). (b) Sanger validation of SLC35G1 in two probands and their parents. (c) Knockout strategy of the mouse *Slc35g1*. Two sgRNAs were designed as indicated. (d) Relative Slc35g1 mRNA level of brain tissues of WT, HET (*Slc35gl*^+/−^), KO (*Slc35gl*^-/-^) mice. (e) Representative heat map for the mouse vs. empty cage trials of the three-chamber test for WT and HET mice. (f) Quantification of social time during (e) of WT and HET mice. (*** *p* < 0.0001, twotailed paired student’ test). (g) Preference index during (e) of WT and HET mice. Calculation methods of preference index see methods (*p* = 0.0018, two-tailed unpaired student’ test). (h) Representative heat map for the familiar partner vs. novel partner trials the three-chamber test for WT and HET mice. (i) Quantification of social time during (h) of WT and HET mice. (** *p* = 0.0022, twotailed paired student’ test). (j) Schematic of the reciprocal social interaction task. Inhabitant: WT or HET, Intruder: WT mice. (k) Quantification of interaction time during (j) of WT and HET mice. (** *p* = 0.0017, two-tailed unpaired student’ test). (l) Representative picture of marble buried after 30 minutes test. (m) Quantification of interaction time during (l) of WT and HET mice. (* *p* = 0.0109, two-tailed unpaired student’ test). Data are the mean±SEM. * *p* < 0.05, \*\**p* < 0.01, \*\*\**p* <0.001

To investigate the functional consequence of *SLC35G1 loss-of-function*, we generated a *Slc35g1* knockout mouse model using two sgRNAs targeting 5’UTR and 3’UTR of the *Slc35g1* gene were used to generate a ~8.5kb deletion (Fig. 5c). We used RT-qPCR to confirm disruption of the *Slc35g1* mRNA from brain tissues collected from WT, *Slc35g1*^+/−^, and *Slc35g1*^-/-^ mice. We found that the mRNA level of *Slc35g1* gene reduced to nearly 50% and 100% in heterozygous and homozygous deletion mice, indicating the success targeting of *Slc35g1* (Fig. 5d).

We next examined *Slc35g1* heterozygous mutant mice (*Slc35g1*^+/−^) with various behavioral tests, which mimicked people with ASD carrying heterozygous mutations of *SLC35G1*. Deficiency of social interaction and stereotypic behaviors are two core symptoms of ASD^1^. We first examined the social behaviors of mice using the classic three-chamber test^21^. We found that *Slc35g1*^+/−^ mice decreased interaction time with partners comparing to wild-type (WT) mice (Fig. 5e, 5f), showing reduced social preference from 35.87% to 5.8% (Fig. 5g). Next, we found that *Slc35g1*^+/−^ mice did not display preference for newly introduced mouse as WT mice did, suggesting that *Slc35g1*^+/−^ mice exhibited defects in both recognizing mice over objects and novel partners (Fig. 5h). To further evaluate the reciprocal social interaction behaviors of *Slc35g1*^+/−^ mice, we performed the intruder interactions test for *Slc35g1*^+/−^ mice (Fig. 5j). Indeed, we found that *Slc35g1*^+/−^ mice exhibited significantly less interaction time with intruder mice, in comparison with WT mice (Fig. 5k). These data strongly indicated that *Slc35g1*^+/−^ mice exhibited deficiency in social interaction behaviors, mimicking the phenotype seen in people with ASD.

We then examined whether *Slc35g1*^+/−^ mice exhibited stereotypic and repetitive behaviors by marble burying test and self-grooming test^21^. We found that *Slc35g1*^+/−^ mice had more frequent marble-burying activity than WT mice (Fig. 5l, 5m), but normal self-grooming activity as WT mice (Fig. S4c), suggesting that *Slc35g1*^+/−^ mice exhibited stereotypic and repetitive behaviors to some extent.

Finally, to examine whether *Slc35g1*^+/−^ mice had abnormal anxiety level, which is commonly associated with people with ASD^21^, we used the open field test and elevated plus maze test. We found that *Slc35g1*^+/−^ mice exhibited a normal level of anxiety level in both tests, comparing to WT mice (Fig. S4d-f). Lastly, we examined the spatial learning ability of *Slc35g1*^+/−^ mice with the Barnes maze test. We found that *Slc35g1*^+/−^ mice showed the same latency in the learning curve as WT mice, in consistent with the finding that human patients carrying *SLC35G1* mutations exhibited normal IQ/DQ (Fig. S4g, h).

It is still a debate if there are “autism-specific genes”. But studying ASD risk genes primarily contributing to social interaction behaviors but not cognitive functions may reveal the neurobiology of social behaviors of mammalian animals. In this study, we analyzed *de novo* variants from ASD with or without DD/ID. We surprisingly found that genes with rare *de novo* variants of large effect (including LGD or severe missense variants) exhibited HFA or LFA-specific patterns. Moreover, our statistical analysis showed that genes with *de novo* variants in LFA are widely associated with various brain regions, whereas genes with *de novo* variants in HFA are enriched in subtypes of neural progenitors (NPC-4), as well as the prenatal birthplace of GABAergic neurons.

Thus, we propose two models for explaining etiology of ASD with or without DD/ID. In the first model, both HFA and LFA genes contribute to the common neural circuits governing social behaviors in mammals, however LFA genes may play additional roles in cognitive development, for example highly expressed in various cortical regions and hippocampus. In the second model, LFA and HFA genes plays distinct role in regulating animal behaviors. For example, the genes with LGD variants in HFA showed enrichment in CGE, MGE, LGE thus likely affecting the proper function of GABAergic neurons, but not affecting the general cognitive function of the brain. We have provided evidence in this study in support of the latter model. In the future, in-depth work illustrating the neural circuits affected by HFA or LFA genes may help to shed light on the neurobiology of social behaviors of mammals.

## Ethics approval and consent to participate

This work is approved by the Ethic Committee of Xinhua hospital, Shanghai Jiao Tong University School of Medicine (XHEC-C-2019-076).

## Availability of data and materials

The datasets used and/or analyzed in the current study are available from the lead contact on the reasonable request.

## Competing interests

The authors declare that they have no competing interests.

## Acknowledgements

The authors thank the ASD families for their participation in this study. We thank technical support team of Mingma Technologies Co., Ltd for whole-exome sequencing analysis. We thank Drs. Aaron Gitler, Yu Fu for providing comments for the manuscript. This work was supported by grants from the NSFC Grants (#31625013, #81941405, #32000726, #82125032, #81930095, #82001211, #81761128035, #31860306); MOST (2019YFA0110100, 2017YFA010330), Shanghai Brain-Intelligence Project from STCSM (16JC1420501); Strategic Priority Research Program of the Chinese Academy of Sciences (XDBS01060200); Program of Shanghai Academic Research Leader, the Open Large Infrastructure Research of Chinese Academy of Sciences, and the Science and Technology Commission of Shanghai Municipality (#2018SHZDZX05, #19410713500 and #2018SHZDZX01), the Shanghai Municipal Commission of Health and Family Planning (GWV-10.1-XK07, 2020CXJQ01, 2018YJRC03 and 2018BR33), and the Guangdong Key Project (2018B030335001), Science and Technology Department of Yunnan Province (202001AV070010), CPSF-CAS Joint Foundation for Excellent Postdoctoral Fellows (2017LH036) and China Postdoctoral Science Foundation (2017M620173).

## Authors’ contributions

All authors contributed to the work and meet the criteria for authorship. Study design: F Li and Z Qiu. Genetic data analysis: J Wang, J Yu, J Wu, K Yang. Acquisition of Clinical information: L Zhang, X Du, F Li. Analysis of single-cell sequencing data: M Wang, X Wang. J Wang performed Sanger verification, RT-PCR, behavior test. Manuscript writing: J Wang, Z Qiu, J Yu. Study supervision: Z Qiu, F Li, X Wang.

## Notes

### Competing Interest Statement

The authors have declared no competing interest.

