## Supplementary material for "*De novo* variants in Chinese ASD trios reveal genetic basis underlying autism without developmental delay and intellectual disabilities": Table S2

| **Brain region** | **abbr.** |
| --- | --- |
| rostral-middle-frontal | RMF |
| pars orbitalis | PAO |
| pars triangulars | PAT |
| superior frontal | SF |
| caudal-middle-frontal | CMF |
| pars opercularis | PO |
| pre-central | PRC |
| frontal lobe | FL |
| occipital lobe | OL |
| parietal lobe | PL |
| temporal lobe | TL |
| post-central | PC |
| supra-maginal | SM |
| superior parietal | SP |
| inferior parietal | IP |
| precuneus | PRECU |
| inferior temporal | IT |
| middle temporal | MT |
| superior temporal | ST |
| bank superior temporal | BST |
| fusiform | FFA |
| lateral occipital | LO |
| insular gyrus | IG |
| medulla | M |
| pons | pons |

**Table S3 Abbreviations for brain regions**
