## Supplemental methods and materials for "*De novo* variants in Chinese ASD trios reveal genetic basis underlying autism without developmental delay and intellectual disabilities"

### Whole-exome sequencing of the Chinese ASD cohort reveals distinct genetic basis underlying autism with or without neurodevelopmental delay

Jincheng Wang[2, #], Juehua Yu[4, #], Mengdi Wang[3, #], Lingli Zhang[1, #], Kan Yang[2, #], Xiujuan Du[1], Jinyu Wu[5], Xiaoqun Wang[3, *], Fei Li[1, *], Zilong Qiu[2, *]

#### Supplemental methods and materials

##### Online database:

1000G-All, 1000G-EAS: https://www.internationalgenome.org/data/

ExAC-All, ExAC-EAS: http://exac.hms.harvard.edu/

SFARI: http://gene.sfari.org

##### Patient Cohort

Unrelated children meeting criteria for ASD according to the Diagnostic and Statistical Manual of Mental Disorders, Fifth Edition (DSM-5) were recruited at Xinhua Hospital from May 2015 to June 2021. Diagnosis was confirmed with Autism Diagnostic Observation Schedule (ADOS) and the Children Autism Rating Scale (CARS) total score of no less than 30. The evaluations were performed by well-trained specialists, psychologists, and developmental behavioral pediatricians. The developmental quotient (DQ) / intelligence quotient (IQ) measure was assessed quantitatively as cognitive ability using age- and development-appropriate standardized tests including the Gesell developmental schedules and Wechsler scales. Patients with neurological disorders (such as cerebral palsy and schizophrenia) or the known chromosome/genetic disorders (such as trisomy 21 syndrome, trisomy 18 syndrome, trisomy 13 syndrome, Rett syndrome, Fragile X syndrome) were excluded from the study.

The study was approved by the ethical committee at Xinhua hospital affiliated to Shanghai Jiao Tong University School of Medicine. Parents or legal guardian of each participant provided written informed consent in compliance with the Declaration of Helsinki.

**Whole-exome sequencing (iGeneTech Bioscience Co., Ltd)**

DNA was extracted from peripheral blood samples with MagPure Tissue&Blood DNA LQ Kit (Magen, Beijing, China) according to the manufacturer's instructions. ~200ng genomic DNA was sheared by Biorupter（Diagenode, Belgium）to acquire 150～200bp fragments. The ends of DNA fragment were repaired and added Illumina Adaptor (Fast Library Prep Kit, iGeneTech, Beijing, China). After sequencing library were constructed, the whole exome were captured with AIExome Enrichment Kit V1 (iGeneTech, Beijing, China) and sequenced on MGI2000 platform with with 150 base paired-end reads.

**Whole-exome sequencing (Mingma Technologies Co., Ltd)**

Peripheral Blood was uniformed into $10^6$ cells/mL, then DNA was extracted using HT Magnetic Purification of DNA from Blood Kit (ZAOTONG) or QIAamp DNA Blood Kit (Qiagen) according to the manufacturer's instructions. In order to test DNA's integrity and concentration, agarose electrophoresis and Qubit 2.0 fluorometer dsDNA HS Assay (Thermo Fisher Scientific) was employed. Finally, high-quality DNA sample (~300ng, $OD_{260/280}$=1.8~2.0) was used for constructing sequencing library.

DNA concentrations were sheared in 150-200bp with Covaris LE220 Sonicator (Covaris). DNA libraries were prepared, repaired and purified with SureselectXT reagent kit (Agilent), End repair mix (component of SureselectXT) and Agencourt AMPure XP Beads (Beckman), respectively. Purified DNA was added with A-tail and ligated with adapter with Sureselect$^{XT}$. Modified DNA fragments were amplified with Herculase II Fusion DNA Polymerase (Agilent). Then exome sequences were captured with SureSelect capture library kit (Agilent). Qubit 2.0 fluorometer dsDNA HS Assay (Thermo Fisher Scientific) was used to measure enriched sequencing libraries. Agilent BioAnalyzer 2100 (Agilent) was used to analyze size distribution of resulting sequencing libraries. According to Illumina-provided protocols for 2x150 paired-end sequencing, paired-end sequencing is performed with Illumina NovaSeq6000 system.

**Variant Calling and Identification of DNMs**

Firstly, we removed adapter bases and low quality bases from raw data, then mapped clean reads to human reference genome (hg19) using BWA[1]. SNVs and indels were called using the sentieon[2] and FreeBayes[3] (version v1.1.0-60-gc15b070, https://github.com/ekg/freebayes) on parent-proband trios. For the mutations called by these two methods, two different filtering criteria were used to identify de novo mutations (DNMs).

Method-1: 1) filter out two or more variant alleles with PASS flag observed in the ExAC population excluding the cohorts of psychiatric disorders (ExAC.r0.3.nonpsych.sites.vcf, N =45,376); 2) filter out one or more variant alleles observed in the 4.7KJPN panel of healthy Japanese individuals released by the Tohoku Medical Megabank Organization of Tohoku University (ToMMo, https://jmorp.megabank.tohoku.ac.jp/dj1/datasets/tommo-4.7kjpn-20190826-af_snvindelall). We used TrioDenovo[4] (version 0.06), a Bayesian framework for detection of DNMs, with the default parameters to extract candidates for DNMs.

Method-2: 1)We used BCFtools (version 1.4.1) to left-align and normalize indels; 2)We retained Variants meeting below criteria: father's genotype was 0/0, the mother's genotype 0/0 and child's genotype either 0/1 or 1/1; father alternate allele count = 0, mother alternate allele count = 0, child allele balance>0.25, father depth > 9, mother depth > 9, child depth> 9, and either child genotype quality (GQ) > 20 (Sentieon) or sum of quality of the alternate observations (QA) > 20 (FreeBayes); 3)We removed variants located in low complexity regions from further analysis (https://raw.githubusercontent.com/lh3/varcmp/master/scripts/LCR-hs37d5.bed.gz). For de novo mutations in chromosome X, we filtered out variants in the pseudoautosomal regions (GRCh37: chrX:60001-2699520 and chrX:154931044-155260560) and the X/Y duplicated transposed region (GRCh37: chrX:88456802-92375509).

Finally, variants obtained through above two filtering frameworks were merged as the final *de novo* mutations.

**Single cell RNA-seq data processing**

Two publics scRNA-seq datasets[5,6] were used to infer the gene expression pattern of

ASD risk genes in developing human brain. R package Seurat[7] was used for downstream analysis. In short, CreateSeuratObject was used to create a Seurat object followed by a Log-normalization process using NormalizeData. Variable genes were identified with FindVariableFeatures.. Principal component analysis was done by RunPCA. Unbiased clustering was performed with FindNeighbors and FindClusters followed by Uniform manifold approximation and projection (UMAP) dimension reduction analysis with function RunUMAP. Differentially expressed genes were selected with FindAllMarkers by setting parameter only.pos =TRUE. Genes with adjusted P-values <0.05.

To study the expression pattern of ASD associate genes, gene-set enrichment analysis was done with R package AUCell[8]. Z-score of the gene-sets enrichment scores were computed with scale and visualized by UMAP.

To explore the expression pattern of four gene-sets among different cell types and brain regions, averaged gene-set enrichment sore was computed and ranked in descending order, and visualized with Heatmap.

To get the proportion of highly expressed genes in four gene-sets(LFA-LGD/LFA-mis/HFA-LGD/HFA-mis), genes numbers expressed in more than 25% cells of each subtypes or brain regions were counted. Total number of genes in each gene-set was divided with highly expressed genes number, the results were considered as proportion of highly expressed genes in each gene set.

**Sanger Sequencing Validation**

The same DNA samples for WES were used for sanger sequencing validation. Primers were designed in the region of 1kb upstream and downstream of mutations to generate not exceeding 800bp PCR products for sanger sequencing. In order to skip special sequence like poly-A or signal attenuation, sequencing primers were designed.

**Genetic modified mice**

All procedures were approved by the Animal Care and Use Committee of the Center for Excellence in Brain Science & Intelligence Technology, Chinese Academy of Sciences, Shanghai, China. Mice were maintained in standard housing conditions on a 12h light/dark cycle(light from 7am to 7pm, dark from 7pm to 7 am) at a constant 22°C with food and water provided ad libitum. Slc35g1 knock-out mice were customized ordered from Biocytogen. Two sgRNAs were designed to generate a 8.5kb chromosomal deletion at Slc35g1 locus in the C57BL/6N genome. Two sequences targeting Slc35g1 were (GCACACGCACCGGCTCGCCG GGG) and (TTCTAGAAGGTTCATCCCGC AGG). Genotypes were determined by PCR of mouse tail DNA, following sequences were used: mutant forward primer (GGCTCCTTGGTTGGGTTACCATTGA) and mutant reverse primer (TGGCTTGTACCTGTGTTCTTTCCGT). Wild-type forward primer (CTGTGCAAAGACAGAGGAGGACCAG), and wild-type reverse primer (TGGCTTGTACCTGTGTTCTTTCCGT).

**Real-time PCR**

In order to quantify mRNA levels, brain RNA was extracted from three pairs littermates of wild-type, slc35g1 heterozygosis and slc35g1 homozygosis mice according to guide of TRIzol™ Reagent. cDNA was reverse transcribed using PrimeScript RT Master Mix (TaKaRa, RR036A). Real-time PCR was carried out with SYBR green premix (Toyobo, QPK-201) and data was analyzed on the StepOnePlus Real-Time PCR System (Applied Biosystems). Gapdh was used as internal control. Primers for Slc35g1:( AACGGGGTTCATAGGTCCCAA) and (CGTTGTCTGGAACGCATAATACA). Primers for Gapdh:( ACGGCCGCATCTTCTTGTGCAGTG) and (GGCCTTGACTGTGCCGTTGAATTT).

**Behavioral assays**

All behavioral assays were performed using 8-12 weeks old male mice. Before behavioral tests, all mice were well handled for 3 days. Mice adapted to testing room at least one hour before tests. Between tests, at least one day rest periods were given. Mice's behaviors were analyzed using Ethovision XT or in a blind manner. For Ethovision analyzing tests, the center point of the mouse was considered as mouse. The equipment used in behavioral tests was cleaned with 75% alcohol to avoiding residual effects of odor.

**Three chambers test**

This test was performed as previously described (Yu et.al 2020 Neurosci Bull.) with minor modifications. Age-matched C57BL/6 male mice were used as social partners. Social partners were placed into wire cage of three-chambers(60cm×40cm×30cm) to habituate for one hour in two days. Test mice habituated three-chambers for 15mins before test.

For sociability test, the partner mouse and empty cage were counterbalanced assigned to left or right chamber. The time spending in partner zone and empty-cage zone was recorded by Ethovision in 10mins. For social memory test, a novel partner was placed into empty cage. The time spending in familiar partner zone and novel partner zone was measured for 10 mins with Ethovision.

**Reciprocal social interactions**

Subjects were housed individually for three days. After ten minutes habituation in testing box, a novel mice was introduced into subject's home cage. Behaviors were recorded for 10 minutes, then manual analyzed. Nose-to-nose sniffing, nose-to-anogenital sniffing, following, pushing past each other with physical contact, crawling over and under each other with physical contact, chasing, mounting and wrestling were regarded as social interactions.

**Marble burying test**

Marble-burying test was performed as previously described (Robert M J Deacon 2006 Nature Protocols). 5cm deep Fresh bedding was filled in testing box(24cm×24cm×24cm). 25 black glass marbles were evenly spaced placed on the surface. Testing mice were placed individually in boxes for 30minutes. Manual count the number of marbles buried.

**Grooming test**

Animals habituated testing box for 20 minutes, then recorded for 10 minutes. Face-wiping, scratching/rubbing of head and ears, and full-body grooming were counted as grooming behavior. The observer manually quantified grooming behavior.

**Elevated plus-maze (EPM) test**

Subjects were placed in the center region of elevated plus-maze, with two open arms(30cm×6cm) and two closed arms(30cm×20cm×6cm), and 50cm above ground. The time spent in open and closed arms was recorded for 10min by Ethovision.

**Open field test**

Open field test was used to test locomotion and anxiety level. Mice was put into an empty, novel 40cm×40cm×40cm square box. Noldus Ethovision analyzed total distance and time spending in the middle of the arena for 10 minutes.

**Barnes maze test**

Strong light was applied for increasing the motivation of mice to hiding in escape box. Training lasted for four days. Mice were placed in square black chamber in the middle of the maze. After ten seconds, chamber was removed. Animal was allowed to explore the maze for three minutes. If it can't find escape box, experimenter guided mice into box. When mice get into escape box, experimenter cover the box for two minutes. Mice received three trials per day for four days. Inter-trial interval was at least 15 minutes. During the training days (day1-day4), the time to find and enter the escape box was manually measured. On the day 5, short-term memory was tested. Mice explored maze for 90s without escape box. Maze was divided into four zones evenly. The time in the zone escape box used was manually measured.
