## Supplemental figures for "*De novo* variants in Chinese ASD trios reveal genetic basis underlying autism without developmental delay and intellectual disabilities"

**Whole-exome sequencing of the Chinese ASD cohort reveals distinct genetic basis underlying autism with or without neurodevelopmental delay**

Jincheng Wang[2, #], Juehua Yu[4, #], Mengdi Wang[3, #], Lingli Zhang[1, #], Kan Yang[2, #], Xiujuan Du[1], Jinyu Wu[5], Xiaoqun Wang[3, *], Fei Li[1, *], Zilong Qiu[2, *]

### Supplemental Figure legends

### Table S1-S3

Figure S1

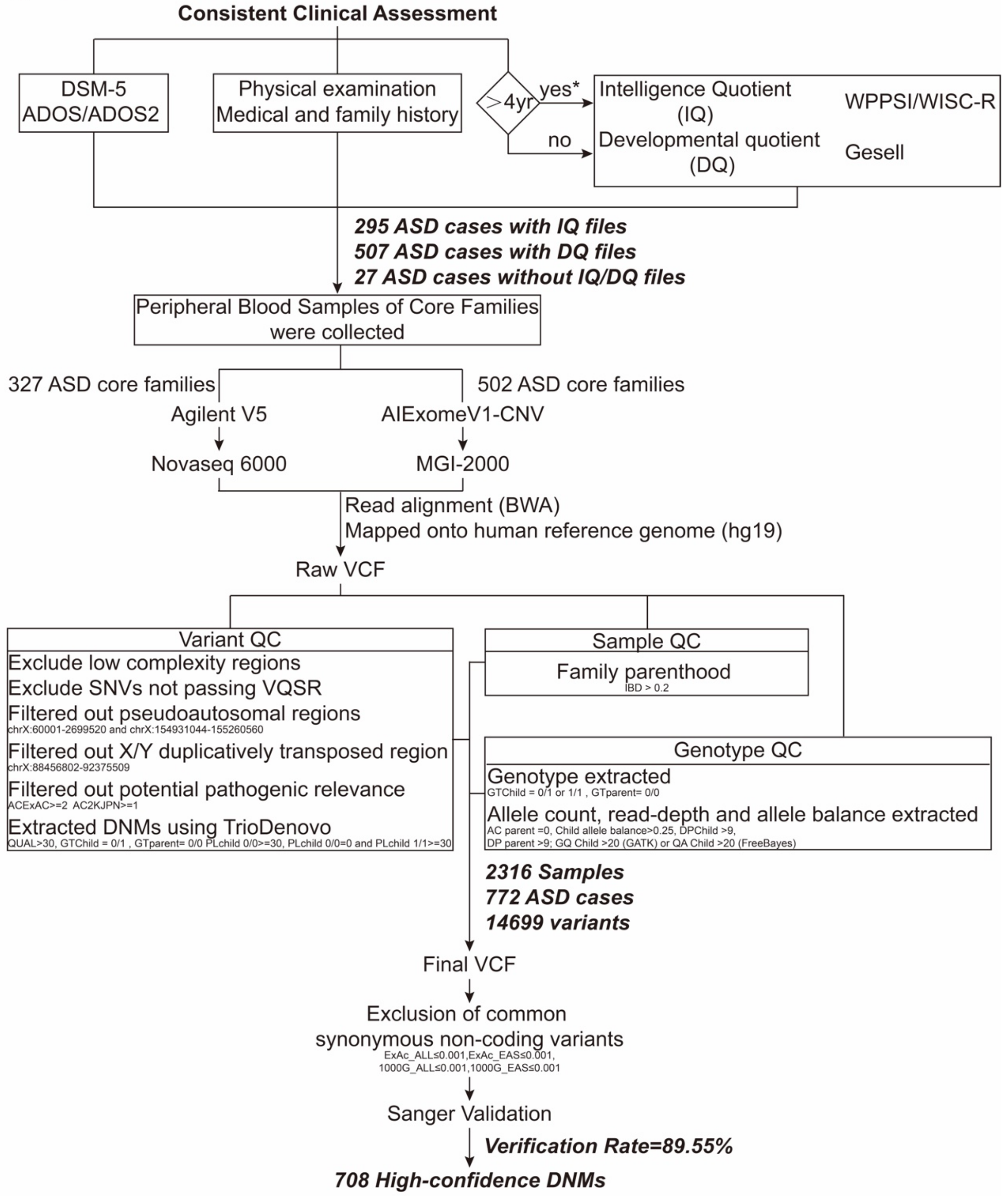

**Supplementary Figure 1. Overview of Clinical Assessment, samples collection and analytical pipeline.**

Children were diagnosed with ASD according to Diagnostic and Statistical Manual of Mental Disorders, Fifth Edition (DSM-5), and were confirmed with Autism Diagnostic Observation Schedule (ADOS) and the Children Autism Rating Scale (CARS). Cognitive ability was measured with developmental quotient (DQ) / intelligence quotient (IQ) according to the Gesell developmental schedules and Wechsler scales. Peripheral blood samples of core families were collected and sequencing with Novaseq-6000 and MGI-2000. Sample quality control steps were applied to the raw joint-genotyped variant calls (VCF) to generate the de novo variants. See Supplemental Experimental Procedures for more detailed descriptions.

Figure S2

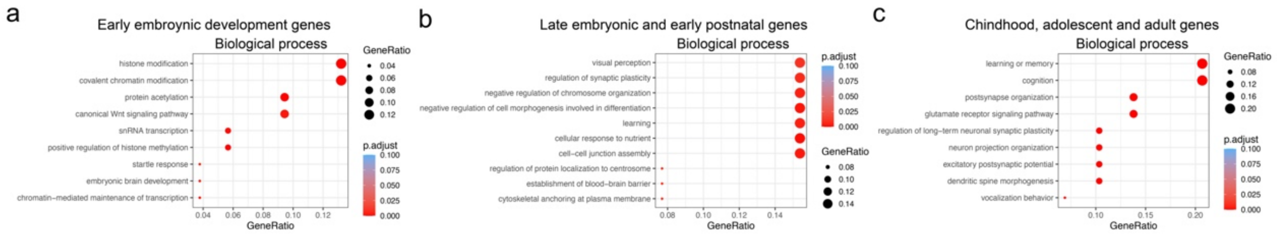


**Supplementary Figure 2 Biological process of gene enrich in different stages.**

Biological process of genes mainly expressed in the early embryonic developmental stages (a), late embryonic and early postnatal stages (b), and childhood, adolescent and adult stages (c).

Figure S3

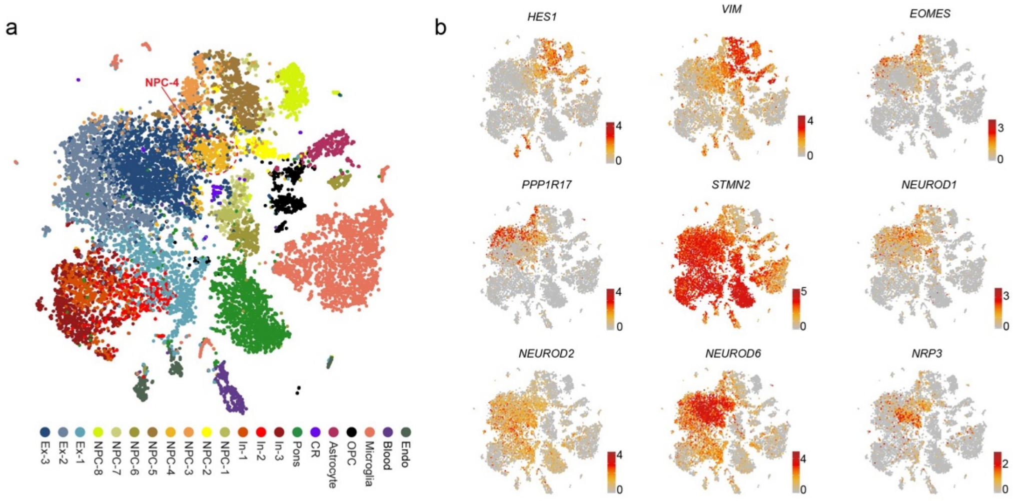


**Supplementary Figure 3 Regional distribution of cells from ASD risk genes enriched subtypes.**

(a) t-SNE showing the clustering of cells from different cortical areas. Cells from cluster NPC-4 were marked with a dotted circle.

(b) Visualization of expression of NSC genes *HES1* and *VIM*, IPC genes *EOMES* and *PPP1R17*, neuronal genes *STMN2* and *NEUROD1/2/6* and layer V gene *NPR3*.

Figure S4

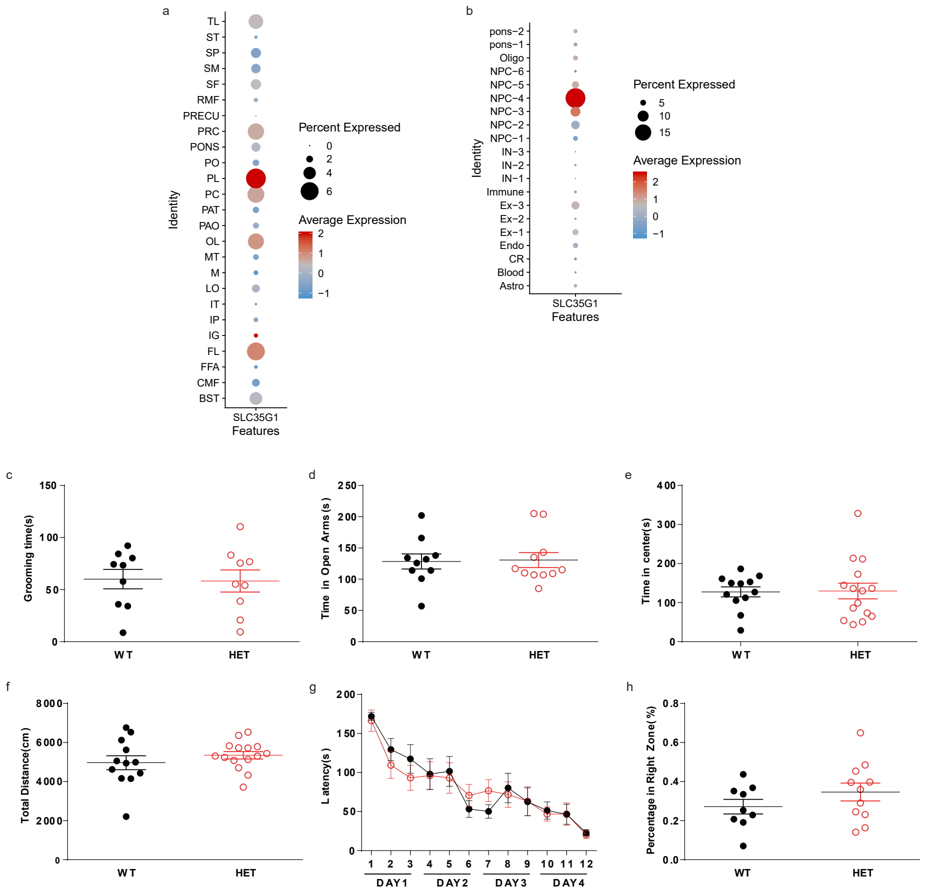


**Supplementary Figure 4 Expression of SLC35G1 in the human brain and behavioral phenotypes of *Slc35g1*+/- mice**

(a) The expression pattern of SLC35G1 in various regions of the human brain.

(b) Expression of SLC35G1 enriched in NPC-4 of the human brain.

(c) Grooming time of *Slc35g1*+/- mice and wild-type mice (two-tailed unpaired t test, $p = 0.8995$).

(d) Time mice spent in the open arm of the EPM test. Two-tailed unpaired t test, $p = 0.8969$.

(e) Time mice spent in the center region of the open field test. Two-tailed unpaired t test, $p = 0.9314$.

(f) The total distance of mice in the locomotion test (two-tailed unpaired t test, $p = 0.3276$).

Latency of mice in the Barnes maze test (g) and percentage in the right zone (h) (g, twoway repeated-measures ANOVA; h, two-tailed unpaired t test, $p = 0.2328$).

**Table S1 List of all *de novo* variants in 772 ASD probands**

**Table S2 Abbreviations of brain regions indicated in Figure 3 and Figure S3**

**Table S3 TADA analysis for all *de novo* variants from 1141 ASD probands**

mut.rate: mutation rate

BF.dn: Bayesian factor for de novo mutation

pval.TADA.dn: TADA p value for de novo mutation

qvalue.dn: q value for de novo mutation (FDR)
